# Kinpute: Using identity by descent to improve genotype imputation

**DOI:** 10.1101/399147

**Authors:** Mark Abney, Aisha El Sherbiny

## Abstract

1

**Motivation:** Genotype imputation, though generally accurate, often results in many genotypes being poorly imputed, particularly in studies where the individuals are not well represented by standard reference panels. When individuals in the study share regions of the genome identical by descent (IBD), it is possible to use this information in combination with a study specific reference panel (SSRP) to improve the imputation results. Kinpute uses IBD information—due to either recent, familial relatedness or distant, unknown ancestors— in conjunction with the output from linkage disequilibrium (LD) based imputation methods to compute more accurate genotype probabilities. Kinpute uses a novel method for IBD imputation, which works even in the absence of a pedigree, and results in substantially improved imputation quality.

**Results:** Given initial estimates of average IBD between subjects in the study sample, Kinpute uses a novel algorithm to select an optimal set of individuals to sequence and use as an SSRP. Kinpute is designed to use as input both this SSRP and the genotype probabilities output from other LD based imputation software, and uses a new method to combine the LD imputed genotype probabilities with IBD configurations to substantially improve imputation. We tested Kinpute on a human population isolate where 98 individuals have been sequenced. In half of this sample, whose sequence data was masked, we used Impute2 to perform LD based imputation and Kinpute was used to obtain higher accuracy genotype probabilities. Measures of imputation accuracy improved significantly, particularly for those genotypes that Impute2 imputed with low certainty.

**Availability:** Kinpute is an open-source and freely available C++ software package that can be downloaded from https://github.com/markabney/Kinpute/releases.

## 2 Introduction

Genotype imputation methods have been a boon to researchers, allowing them to maximize available resources by allowing them to computationally infer the alleles at untyped variants for many individuals. Although a majority of genotypes may be imputed with high accuracy using standard reference panels, a substantial fraction of variants will typically be discarded due to low quality imputation. For study populations that are not well represented by available reference panels, this problem is exacerbated (Herzig *et al*., 2018). When imputing genotypes in indigenous and founder populations, or even isolated populations of European ancestry, the results can be significantly improved when a subset of the sample are sequenced and included as a study specific reference panel (SSRP) (Deelen *et al*., 2014; Hou *et al*., 2017; Mitt *et al*., 2017; Pistis *et al*., 2015; Sidore *et al*., 2015; Zhou *et al*., 2017; Herzig *et al*., 2018).

When identity by descent (IBD) from close relatives to an unsequenced individual exists, linkage disequilibrium (LD) based imputation approaches are expected to have high accuracy (McCarthy *et al*., 2016). This is because LD-based approaches will, in principle, phase and match the haplotype of the unsequenced individual with the IBD segment from a sequenced individual. In practice, this phasing and matching are done imperfectly, allowing for the possibility that IBD based imputation methods might be an alternative. IBD based imputation methods generally are either ones that find and use extended IBD regions to both phase and impute genotypes (Kong *et al*., 2008; Palin *et al*., 2011; Uricchio *et al*., 2012) or use a pedigree to inform imputations (Burdick *et al*., 2006; Chen and Schaid, 2014; Cheung *et al*., 2013; Livne *et al*., 2015). Pedigree based imputation can outperform LD based methods, particularly at variants with low allele frequencies (Ullah *et al*., 2019), but it is less clear whether IBD can improve on LD based imputation in the absence of a pedigree once an SSRP is included (Herzig *et al*., 2018).

Here, we show that our method, Kinpute, which does not use pedigree data, can use IBD information to obtain substantially improved imputation accuracy. Kinpute uses a novel approach to imputation in that genotype probabilities output from LD imputation methods, such as Impute2 (Howie *et al*., 2012), minimac (Fuchsberger *et al*., 2015; Das *et al*., 2016), and Beagle (Browning *et al*., 2018; Browning and Browning, 2016) are used as prior probabilities in a unified probabilistic model that includes IBD information. Other approaches to combine IBD and LD based imputation include (1) selecting either the pedigree imputed genotype, if a pedigree based method can be used, or the LD imputed genotype according to some criterion (Saad and Wijsman, 2014; Blue *et al*., 2014; Livne *et al*., 2015) or (2) by taking a linear combination of the two sets of imputed genotypes (Chen and Schaid, 2014). By using the LD imputed genotypes as prior probabilities, Kinpute provides a natural, probabilistic model to integrate the two sources of imputation information while allowing for a wide range of study designs (e.g. presence or absence of pedigree data). In addition, Kinpute does not require phase information, making it robust to phasing errors, but does require an SSRP, which may also be used when performing LD-based imputation, and implements a novel algorithm to select an optimal SSRP.

## 3 Methods

The software consists of two components, both of which require prior estimates of IBD sharing. The first component selects an optimal set of individuals to use as an SSRP while leaving the rest as the imputation panel. The second component performs genotype imputation on the imputation panel given sequence data on the SSRP, prior genotype probabilities, and the previously computed IBD information.

### 3.1 Optimal study specific reference panel

To select an optimal SSRP we assume knowledge of kinship coefficients, computed from a pedigree or from a genetic relationship matrix estimated from genotype data, for every pair of individuals in the study. These coefficients measure the probability of a pair of individuals being IBD at an arbitrary locus.

We seek a set of individuals of a prespecified size from the study sample that, when sequenced, will provide the most informative subsample (i.e. the SSRP) to impute the sequence data into the remaining individuals (i.e. the imputation sample). We use IBD sharing as the measure of informativeness for imputation. Consider a locus where IBD probabilities have been computed between all pairs in the sample. Let {*ϕ_ij_*} be the set such that *ϕ_ij_* is the probability that, at that locus, an allele drawn randomly from individual *i* is IBD with a randomly drawn allele from individual *j*. When computed from a pedigree in the absence of marker data, this definition is formally equivalent to the kinship coefficient. Also, let *R* be the set of sequenced individuals and *U* the set of genotyped but not sequenced individuals, and let *r* ∈ *R* and *u* ∈ *U*. The quantity *ϕ_ru_*, then, is a measure of how informative *r* is with respect to imputing *u*’s genotype at that locus. For two sequenced individuals *r*_1_, *r*_2_ our information for imputing *u*’s genotype is measured by 1 – (1 – *ϕ*_*r*_1_*u*_)(1 – *ϕ*_*r*_2_*u*_). This is the probability that at least one of the randomly drawn alleles from the sequenced individuals will be IBD with a random allele from *u* at the locus. As this probability approaches one we become increasingly certain that at least one of the sequenced individual will be IBD with *u*. We extend this pattern to form a score by taking the product over all *r* ∈ *R* and summing over all *u* ∈ *U* for a given division of subjects into *R* and *U*. The optimal set of individuals to sequence at this locus is obtained by minimizing the loss function 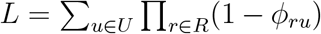 over all divisions of subjects for a given number of subjects in *R*. To obtain a genomewide score we let *ϕ_ij_* be either the kinship coefficient or the *ij*th component of the genetic relatedness matrix computed from known genotype data. Alternatively, one may define a value *L_m_* at each marker *m* and set 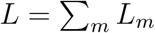, though we do not implement this latter approach.

We use a simple greedy algorithm to select which individuals in the sample to include in *R*. That is, the first individual from the sample is chosen such that *L* is minimized for that person in *R* and everyone else in *U*. Subsequent individuals are added to *R* by keeping the individuals already in *R* and finding which one of the remaining individuals in *U* minimizes *L*. This procedure is continued until the desired size of *R* is met. Note that this algorithm will not, in general, result in the globally optimal solution for *R*. It does, however, provide a computationally tractable approach to obtaining a good, if not the best, solution.

Note that under the case where we are perfectly informed with respect to IBD sharing, the method proposed in (Gusev *et al*., 2012) is a special case of our more general algorithm. In their approach, rather than basing the loss function on IBD probabilities it is based on hard calls of segments being IBD or not. IBD segment calling, however, is in practice far from perfect and can have significant false positive and/or false negative error rates. Using probabilities has the advantage of taking uncertainty into account and allowing moderate probabilities to combine for greater information. This is not possible if the probabilities are thresholded to a binary classification of IBD or not.

### 3.2 Imputation

The imputation method is designed to work in conjunction with the output of other genotype imputation methods, e.g. Impute2 (Howie *et al*., 2012), minimac (Fuchsberger *et al*., 2015) or Beagle (Browning and Browning, 2016), that output genotype probabilities at specific markers. These genotype probabilities are used as prior probabilities for our method. In the absence of genotype probabilities from other methods, Kinpute will compute prior probabilities from the allele frequencies in the SSRP.

Consider a study where the entire sample has been genotyped at a set of markers (i.e. the framework markers) and in which a subset of the sample of individuals have been sequenced. The sequenced individuals are the SSRP *R* = {*r_i_*}, where *r_i_* is the ith reference panel individual, *i* = 1,…, *N*. Let 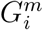 be the true, unphased sequence data at marker *m, m* = 1,…, *M* in the SSRP with *i* = 1,…, *N*, and 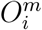 be the corresponding observed, unphased sequence data. The imputation problem is to find 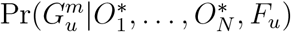, where *F_u_* is the framework genotype data of *u*, 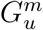 is the unknown sequence at marker m of sample individual u who has not been sequenced and the superscript asterisk indicates the set over all markers. In addition, we assume that from the framework genotype data we have estimates 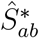 on the IBD condensed identity states 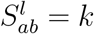 where *k* ∈ {1,…, 9} indexes the identity state at framework marker *l, l* = 1,…, *L* < *M*, and *a, b* ∈ {*u, r*_1_,…, *r_N_*}. (Supplementary Figure 1 displays the nine condensed identity states.)

Standard imputation methods typically use LD information to first phase the observed genotype and sequence data then compute the probability 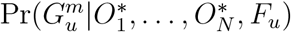. That is, a primary component of the imputed genotype at a marker is the phased observed alleles of u at nearby markers. Errors in phasing, or in the observed genotypes or sequences, can reduce the accuracy of the imputation. Kinpute on the other hand, uses IBD, which extends over a longer range than LD. Given this IBD information Kinpute seeks to compute the probability 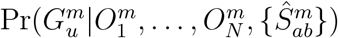. Note that in LD-based imputation the probability is computed given the genomewide sequence data of the reference panel and genomewide framework genotype data, while the conditional probability used by Kinpute apparently conditions only on the reference panel sequence data and IBD estimates at marker *m*. In fact, Kinpute implicitly conditions on the genomewide framework data because this is used in the computation of the IBD estimates 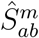 and on the reference panel genomewide sequence through the use of a prior genotype probability that depends on this, as described below. Note also that though the probabilities are conditioned on genomewide data, in practice information local to m is most influential. Rather than compute this conditional probability directly under some probability model (often some form of a hidden Markov model), as is done with LD based imputation, we instead use Bayes’ law,

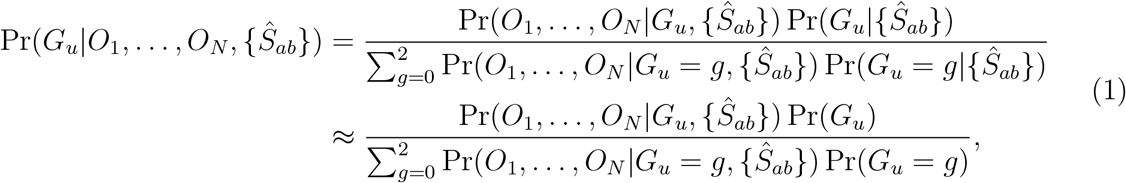

where we have dropped the marker *m* superscript for notational simplicity. We have used Pr(*G_u_* = *g*|{*Ŝ_ab_*}) ≈ Pr(*G_u_* = *g*), which is an approximation because, for instance, the presence of inbreeding can affect the probabilities of *u*’s genotypes. Unless inbreeding is substantial, however, the approximation is highly accurate. Furthermore, this formulation allows us to choose prior probabilities Pr(*G_u_*) based on other sources of information. In particular, we use the imputed genotype probabilities of standard LD based imputation as prior probabilities. Below, we focus on our approach to computing Pr(*O*_1_,…, *O_N_*|*G_u_*, {*Ŝ_ab_*}).

#### 3.2.1 Error model

Throughout, we allow for errors to exist between the true genotype *G_i_* and the observed genotype *O_i_*. We further assume that the observed genotypes are conditionally independent of all other random variables given the true genotypes so that, for instance, Pr(*O*_1_, *O*_2_|*G*_1_, *G*_2_) = Pr(*O*_1_|*G*_1_) Pr(*O*_2_|*G*_2_). We assume an allelic error rate of e with errors of the two alleles in a genotype being independent and the probability of error at different markers also independent. (See Supplementary Table 1). This error model is certainly a simplification; however, it does not play a strong role in the final imputed genotype probabilities. Primarily, the error model prevents numerical failure of the algorithm that can occur from inconsistent observations, for instance, when the observed genotypes are inconsistent with the IBD state. In our tests small non-zero error rates, up to at least 5%, had only minor impact on the imputed probabilities. In the Kinpute software package we set *ϵ* = 0.005.

#### 3.2.2 Independence within the reference panel

At times, one simplifying assumption we use is that the genotypes of the reference panel individuals are conditionally independent given *G_u_*. In this case we have

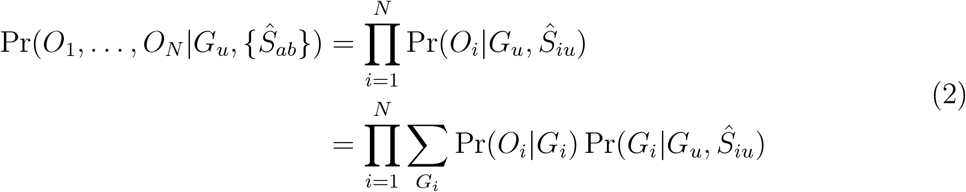

The estimated IBD information *Ŝ_iu_* consists of probabilities of each identity state given the framework set of markers, 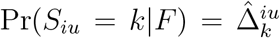, where *k* = 1,…, 9 indexes the identity state. That is, 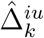 is the estimated locus-specific probability of the identity state given the genotype data, and we assume this is already known. In practice, we compute these probabilities using the IBDLD (Han and Abney, 2011, 2013) software package. Furthermore, we have

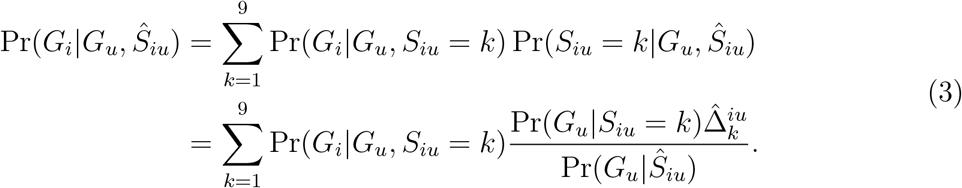

In the above we use

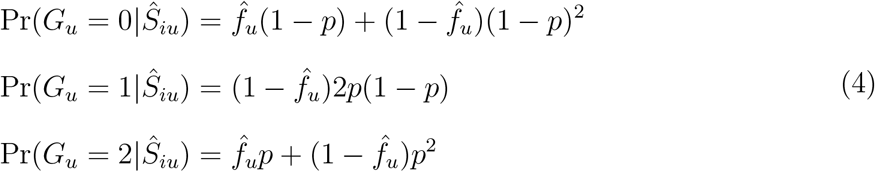

where 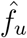 is the estimated inbreeding coefficient of individual *u* at the current marker, and *p* is the frequency of the 1 allele. The probabilities Pr(*G_u_*|*S_iu_* = *k*) are given in Supplementary Table 2, and the probabilities Pr(*G_i_*|*G_u_, S_iu_* = *k*) are given in Supplementary Table 3. The quantities in these two tables together with equation (3) allows us to compute the probability in equation (2).

#### 3.2.3 IBD within the reference panel

It may often be the case that IBD exists between members of the reference panel at some markers. We can leverage this IBD to obtain more accurate estimates of the genotype probability of individual u at that marker. We do this by altering the factoring that occurs in Equation (2) to reflect this dependence. For instance, let the reference panel be composed of three individuals 1, 2, 3 and there is an individual u in whom we are imputing the genotypes at a SNP. Assume that individuals *u*, 1, 2 are in an informative 3-way IBD state *S*_*u*12_ with some probability Ψ_*u*12_, while individual 3 is not IBD with either 1 or 2. We get,

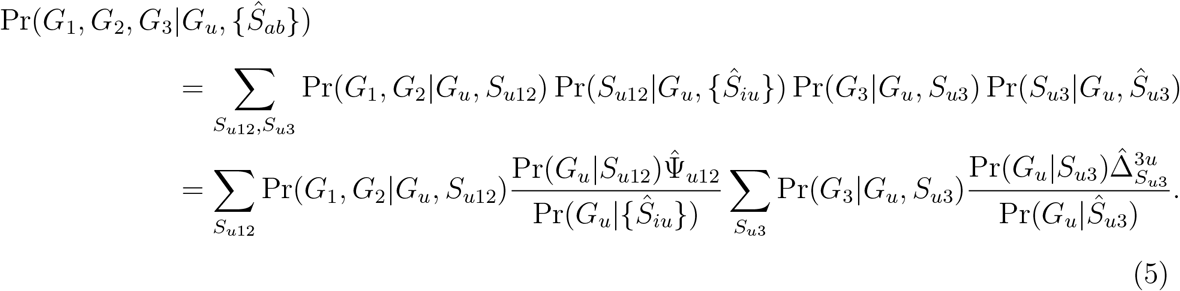

The probabilities Pr(*G_u_*|*Ŝ_iu_*) are given by equations (4). There are many possible 3-way IBD states at a locus (Thompson, 1974), and computing them would be challenging. Instead, we use an approximation based on the pairwise IBD states at the locus and define *S*_*u*12_ = (*S*_*u*1_, *S*_*u*2_, *S*_12_). Then, letting *s* = (*S*_*u*1_ = *k, S*_*u*2_ = *l, S*_12_ = *m*) such that (*k, l, m*) define a legal 3-way IBD state, we define the probability of the 3-way IBD configuration as,

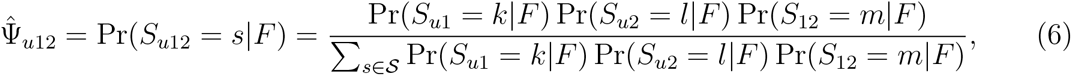

where 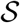 is the set of only the eight most probable values of *s*, for computational expediency.

In a collection of reference panel individuals, we need an algorithm to place them into 3-way configurations with *u*. Random, or arbitrary groupings are unlikely to be generally useful as only some 3-way configurations will be informative for imputation.

The algorithm proceeds by first picking an individual u into whom we wish to impute genotypes. At each marker, the reference panel individuals are grouped into 3-way configurations (as listed in Supplementary Table 4), if possible. Reference panel individuals who do not end up in a 3-way configuration are treated as independent. For purposes of classifying into configurations, the IBD state for a pair (e.g. *S*_*u*1_) is the IBD state with the highest probability. If a reference panel individual belongs to more than one configuration, only one of them is chosen. For computational expediency we do not attempt to optimize the placement into configurations (some configurations may be more informative than others based on the genotypes of the individuals), and instead simply place a reference panel individual into the first configuration found. Once reference panel individuals are placed into configurations the genotype probabilities are computed as given in Supplementary Tables 5 – 13, individuals who are not in a configuration have genotype probabilities given by Supplementary Table 3. These probabilities are combined, as in Equation (5), with 3-way configurations weighted according to their probability, as computed by Equation (6). We further apply our error model to obtain,

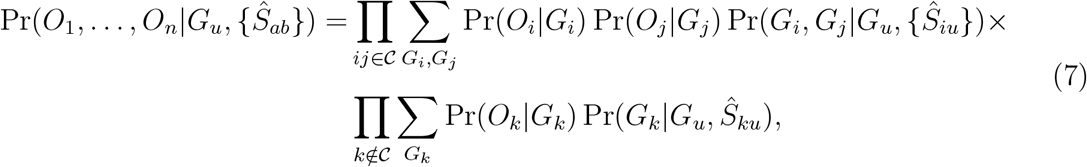

where 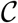 is the set of all reference panel individuals in a 3-way configuration. The final imputed genotype probabilities are, given by Equation (1).

### 3.3 Data and analysis

We applied our method to a human founder population. The study comprised 1,415 individuals from the Hutterite kindred. A framework set of markers, consisting of 271,486 SNPs that passed quality control and had a minor allele frequency of at least 0.05, was obtained from a combination of three Affymetrix arrays (500k, 5.0 and 6.0). The framework markers were not pruned for LD. From these individuals, 98 were sequenced by Complete Genomics, resulting in 6,617,627 SNPs that passed quality control. Details of the genotype and sequence data, including quality control is described in Livne *et al*. (2015). Although these individuals are linked by a known pedigree, all our analyses described below are done without knowledge of the pedigree.

From the 98 sequenced individuals, 50 were chosen to act as an SSRP while the other 48 were used as subjects for imputation given their framework genotypes. The sequence data of the 48 were hidden from imputation and used as the “ground truth” against which to compare the imputed genotypes. To assess the degree to which Kinpute can use IBD information to improve genotype imputation we focus on the genotypes on chromosome 22 only. On chromosome 22 there are 3059 SNPs in the framework set and 111,369 SNPs in the 98 sequenced individuals. We performed an initial round of imputation on the 48 subjects using Impute2 (Howie *et al*., 2012) with pre-phasing by ShapeIt2 (O’Connell *et al*., 2014). We used a reference panel that consisted of the 1000 Genomes panel (1000 Genomes Project Consortium *et al*., 2015) merged with the 50 SSRP individuals, with merging performed by Impute2. The resulting imputed genotype probabilities were used as prior probabilities for our Kinpute method, as described below.

To apply our Kinpute method we first used the IBDLD software package (Han and Abney, 2011) with the GIBDLD method (Han and Abney, 2013), which does not use a pedigree, to estimate probabilities of IBD sharing at each SNP in the framework set for every pair of the 98 individuals. Using these IBD probabilities, the SSRP, and the output from Impute2 as the genotype prior probabilities, we used Kinpute to obtain posterior probabilities for every genotype on chromosome 22 in the 48 subjects. We then compared the imputed genotypes, both those done solely by Impute2 and those done with the combination of Impute2 and Kinpute, with the sequenced genotypes in the 48 subjects to assess accuracy.

## 4 Results

To assess the imputation improvement provided by Kinpute, we stratify the imputed genotypes by genotype and SNP metrics. SNPs are stratified by Impute2 Info score (> 0.4 or < 0.4), minor allele frequency (≤ 0.02 or > 0.02), and whether the SNP was shared (i.e. in both reference panels), or private (i.e. only in the SSRP). Minor allele frequency was computed from the pooled reference panels for shared SNPs, or from the SSRP for private SNPs. In addition, each imputed genotype was stratified based on whether the Impute2 results had high certainty (i.e. the maximum probability for a genotype was at least 0.9), or low certainty (i.e. the complement of high certainty). Within each stratification bin, we compute imputation quality metrics using all genotypes in that bin. In Table 1 we show the genotype dosage *R*^2^ between the true genotype and the imputed dosages computed from Impute2 and from Kinpute with Impute2 priors. The additional quality metrics, heterozygote sensitivity, heterozygote positive predictive value, and concordance are given in Supplementary Tables 14-16.

**Table 1:** Imputed genotype dosage *R*^2^ with the true genotypes

| Info | SNP type | MAF | genotype<br>certainty | Number of<br>genotypes (%) | Impute2 | Impute2 &<br>Kinpute |
| --- | --- | --- | --- | --- | --- | --- |
| > 0.4 | shared | > 0.02 | High | 2911525 (59.7) | 0.90 | **0.93** |
| > 0.4 | shared | > 0.02 | Low | 469987 (9.6) | 0.32 | **0.68** |
| > 0.4 | shared | ≤ 0.02 | High | 387932 (7.9) | 0.90 | **0.94** |
| > 0.4 | shared | ≤ 0.02 | Low | 15460 (0.3) | 0.25 | **0.70** |
| > 0.4 | private | > 0.02 | High | 201672 (4.1) | 0.89 | **0.92** |
| > 0.4 | private | > 0.02 | Low | 31896 (0.7) | 0.35 | **0.66** |
| > 0.4 | private | ≤ 0.02 | High | 54182 (1.1) | 0.53 | **0.66** |
| > 0.4 | private | ≤ 0.02 | Low | 3178 (0.1) | 0.07 | **0.47** |
| < 0.4 | shared | > 0.02 | High | 235481 (4.8) | 0.63 | **0.80** |
| < 0.4 | shared | > 0.02 | Low | 106855 (2.2) | 0.28 | **0.73** |
| < 0.4 | shared | ≤ 0.02 | High | 284529 (5.8) | 0.61 | **0.77** |
| < 0.4 | shared | ≤ 0.02 | Low | 8751 (0.2) | **0.13** | 0.0 |
| < 0.4 | private | > 0.02 | High | 28684 (0.6) | 0.63 | **0.82** |
| < 0.4 | private | > 0.02 | Low | 7988 (0.2) | 0.34 | **0.76** |
| < 0.4 | private | ≤ 0.02 | High | 123111 (2.5) | 0.45 | **0.52** |
| < 0.4 | private | ≤ 0.02 | Low | 5913 (0.1) | 0.06 | **0.36** |

In all but one stratification level, where neither method does well, Kinpute improves the imputation quality of Impute2. For genotypes that are of low certainty, in particular, Kinpute can substantially increase *R*^2^, especially for SNPs that have low INFO score.

## 5 Discussion

We have shown that in the presence of IBD, even when pedigree data is absent, Kinpute can significantly improve the quality of imputed sequence data over using LD based imputation methods alone. Note that Kinpute should be viewed as a supplement, as opposed to an alternative, to LD-based imputation methods such as Impute2, Beagle or minimac. These LD-based methods provide useful information to Kinpute’s IBD-based approach in the form of prior genotype probabilities. The IBD approach used by Kinpute is distinct from pedigree-based approaches, which also use IBD. Pedigree approaches (Burdick *et al*., 2006; Chen and Schaid, 2014; Cheung *et al*., 2013; Livne *et al*., 2015) require the pedigree to estimate IBD, often via the construction of inheritance vectors. Thus only sequence data within the pedigree can be used for the IBD-based imputation. When a pedigree has only one or a few individuals who were sequenced, pedigree based imputation will be of limited use. For instance, in the Hutterite data set used here, we were able to assess the imputation accuracy by holding out 48 sequenced individuals while using 50 sequenced individuals as an SSRP. Even though a pedigree is available in the Hutterite population, the subpedigrees that would have been needed for the pedigree based methods (Burdick *et al*., 2006; Chen and Schaid, 2014; Cheung *et al*., 2013) would have left too few SSRP individuals in any subpedigree for the methods to have effectively imputed genotypes in the imputation panel. Kinpute, on the other hand, does not need a pedigree and can use IBD from all sequenced individuals, including those that are cryptically related. This is particularly useful in populations where many individuals may be related, but do not have a recorded pedigree.

Kinpute is advantageous in that it can use IBD even from distant or cryptically related individuals to improve LD-based imputation results. However, like any IBD based method, its utility is limited by the amount of IBD that is detectable in the sample. For samples that come from large, outbred populations, where IBD is sparse, every IBD-based method will be of limited use. When imputing a genotype where the information from IBD goes to zero, the genotype probabilities returned from Kinpute will converge to the provided prior probabilities, typically the LD-based estimates. As samples increase in IBD, Kinpute’s performance will improve. Samples from populations such as the Hutterites, where IBD is common, will gain the most. Similar types of gains in imputation can be expected from studies on population isolates, indigenous groups, and endogenous populations. Note that having as high a level of IBD as in the Hutterites is not required to gain from using Kinpute. Much of the imputation gain results from the novel use of IBD configurations. For instance, though an IBD value of 1 between an unknown genotype and a SSRP genotype provides only limited information for imputation, when this IBD occurs within certain configurations, the information for imputation can be dramatically higher. One instance where this arises is when the SSRP individual’s genotype is heterozygous (i.e. (0, 1)) and is also IBD = 1 with a homozygous (e.g. (0, 0)) SSRP genotype. If the unknown genotype has an IBD value of 0 with this second SSRP genotype, we can immediately conclude that the unknown genotype must have at least a 1 allele (Figure 1). When the 1 allele is rare, LD based imputation often has trouble imputing the presence of this allele in the unknown genotype. In this situation, even though the amount of IBD is not high, IBD provides significant added value and results in both much higher heterozygote sensitivity (Table 14 in the Supplementary Information) and heterozygote positive predictive value (Table 15 in the Supplementary Information).

**Figure 1:**
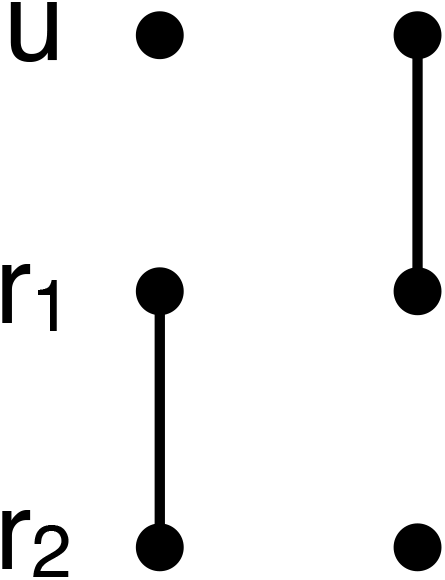
Highly informative IBD configuration. Individual u has IBD 1 with *r*_1_ and IBD 0 with *r*_2_ while *r*_1_ and *r*_2_ have IBD 1. When *r*_1_ has genotype (0, 1) and *r*_2_ has genotype (0, 0), we can immediately infer that *u* must have at least one copy of the 1 allele.

Kinpute relies on the results of other computational tools and, as a consequence, the final results can be dependent on the tools used and their accuracy. When selecting an optimal sample to sequence, Kinpute requires kinship coefficients. These values may differ depending on whether the coefficients are computed from a pedigree or from genotype data, and, when computed from genotype data, what method was used. Different estimates of the kinship coefficient may lead to different samples chosen for the SSRP but do not directly affect the imputation. Insofar as different methods give similar estimates of the kinship coefficients, we expect the selected SSRPs will be similar and the imputation results to be largely unaffected. When performing imputation, Kinpute requires prior probabilities for each genotype. We recommend that the output of LD-based imputation be used, but any set of prior probabilities are allowed. If no prior probabilities are provided, Kinpute will use allele frequencies in the SSRP to compute prior genotype probabilities assuming Hardy-Weinberg equilibrium. Depending on how informative the IBD is at a genotype, the posterior imputed probabilities may be sensitive to the priors used.

As a general rule, the larger the SSRP, the more accurate LD based imputation will be. Similarly, when a sample comes from a population that is close to a population in the standard reference panels, LD based imputation will generally be of high accuracy. However, even in this case, with a reference panel in the tens of thousands, some genotypes, particularly at SNPs with a rare minor allele frequency, are not very well imputed (McCarthy *et al*., 2016). Approaches that can measurably improve imputation quality, then, can be of great benefit. This is particularly true when budgets limit the number of individuals that can be sequenced and the study population has drifted significantly from those in standard reference panels. When relatedness exists in the sample, whether close, distant or cryptic, Kinpute provides an additional useful tool to maximize the use of the study’s sequence data.

## 7 Acknowledgements

We thank Dr. Carole Ober for use of the Hutterite data.

