## Supplemental information for "Kinpute: Using identity by descent to improve genotype imputation"

4 Informative 3-way configurations. Note that because individuals 1 and 2 are unordered, we list only one of the corresponding ordered 3-way configurations (e.g. configuration (8, 9, 8) is also informative but has the same probabilities as configuration (9, 8, 8) with individuals 1 and 2 switched). Note that for configuration (8, 8, 7), (7, 7, 7) and (3, 3, 7) $\Pr(G_1, G_2|G_u, S_{u12}) = \Pr(G_1|G_u, S_{u1}) = \Pr(G_2|G_u, S_{u2})$, though we do not seek these configurations in our algorithm. . . . . . 7

5 Configuration ($S_{u1} = 9, S_{u2} = 8, S_{12} = 8$), probabilities $\Pr(G_1, G_2|G_u, S_{u12})$ . 7

6 Configuration ($S_{u1} = 8, S_{u2} = 8, S_{12} = 9$), probabilities $\Pr(G_1, G_2|G_u, S_{u12})$ . 7

7 Configuration ($S_{u1} = 8, S_{u2} = 8, S_{12} = 8$), probabilities $\Pr(G_1, G_2|G_u, S_{u12})$ . 8

8 Configuration ($S_{u1} = 8, S_{u2} = 6, S_{12} = 5$), probabilities $\Pr(G_1, G_2|G_u, S_{u12})$ . 8

9 Configuration ($S_{u1} = 8, S_{u2} = 5, S_{12} = 6$), probabilities $\Pr(G_1, G_2|G_u, S_{u12})$ . 8

10 Configuration ($S_{u1} = 8, S_{u2} = 5, S_{12} = 5$), probabilities $\Pr(G_1, G_2|G_u, S_{u12})$ . 9

11 Configuration ($S_{u1} = 5, S_{u2} = 5, S_{12} = 2$), probabilities $\Pr(G_1, G_2|G_u, S_{u12})$ . 9

12 Configuration ($S_{u1} = 4, S_{u2} = 3, S_{12} = 8$), probabilities $\Pr(G_1, G_2|G_u, S_{u12})$ . 9

13 Configuration ($S_{u1} = 3, S_{u2} = 2, S_{12} = 5$), probabilities $\Pr(G_1, G_2|G_u, S_{u12})$ . 10

14 Heterozygote sensitivity . . . . . 10

15 Heterozygote positive predictive value . . . . . 11

16 Concordance rates . . . . . 12

### List of Figures

1 The nine condensed identity states $S_{AB}$. A line joining two alleles indicates those alleles are IBD. . . . . . . . . . . . . . . . . . . . . . . . . . . . . . . . 5

### 1 Additional quality metrics

In addition to $R^2$, we computed other measures on how much Kinpute improves the imputation quality of LD-based methods. In all measures we compare the imputed genotypes (both from Impute2 and from Kinpute) to the true genotypes on the 48 held out individuals. Genotypes were compared only for chromosome 22. The additional measures are, concordance, heterozygote sensitivity (i.e. recall), and heterozygote positive predictive value (i.e. precision). Each of these measures require hard genotype calls, which were done at a genotype probability threshold of 0.9, (i.e. genotypes classified as having "high" certainty) or by picking the genotype with the maximum probability (i.e. for genotypes classified as having "low" certainty). The concordance is defined as the frequency that the called genotype is the same as the true genotype. Heterozygote sensitivity is the fraction of true heterozygotes that are called as heterozygotes. The heterozygote positive predictive value is the fraction of called heterozygotes that are true heterozygotes. Results are shown in Tables 14 – 16.

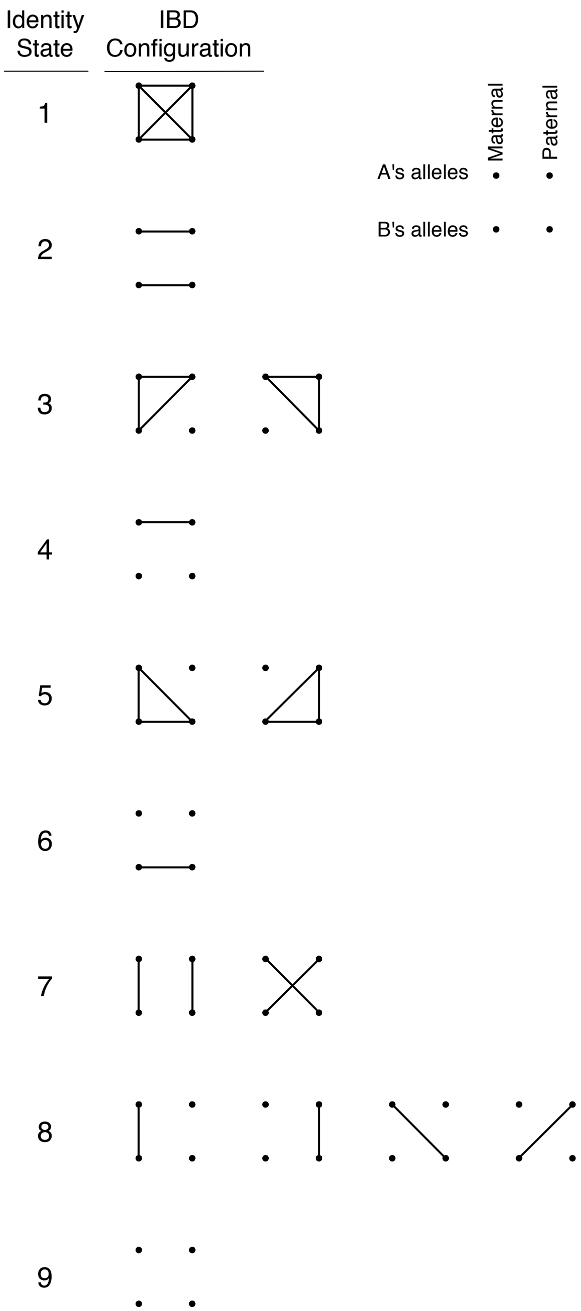

Figure 1: The nine condensed identity states $S_{AB}$. A line joining two alleles indicates those alleles are IBD.

| | $O_i$ | | |
|---|---|---|---|
| $G_i$ | 0 | 1 | 2 |
| 0 | $(1-\epsilon)^2$ | $2\epsilon(1-\epsilon)$ | $\epsilon^2$ |
| 1 | $\epsilon(1-\epsilon)$ | $1-2\epsilon(1-\epsilon)$ | $\epsilon(1-\epsilon)$ |
| 2 | $\epsilon^2$ | $2\epsilon(1-\epsilon)$ | $(1-\epsilon)^2$ |

Table 1: Genotype error model, $\Pr(O_i|G_i)$

| | $G_u$ | | |
|---|---|---|---|
| $k$ | 0 | 1 | 2 |
| $1,2$ | $(1-p)$ | 0 | $p$ |
| $3,4$ | $(1-p)^2$ | $2p(1-p)$ | $p^2$ |
| $5,6$ | $(1-p)$ | 0 | $p$ |
| $7,8,9$ | $(1-p)^2$ | $2p(1-p)$ | $p^2$ |

Table 2: $\Pr(G_u|S_{iu}=k)$

| | | $G_i$ | | |
|---|---|---|---|---|
| | $G_u$ | 0 | 1 | 2 |
| $S_{iu}=9$ | $0,1,2$ | $(1-p)^2$ | $2p(1-p)$ | $p^2$ |
| $S_{iu}=8$ | 0 | $1-p$ | $p$ | 0 |
| | 1 | $(1-p)/2$ | $1/2$ | $p/2$ |
| | 2 | 0 | $1-p$ | $p$ |
| $S_{iu}=7$ | 0 | 1 | 0 | 0 |
| | 1 | 0 | 1 | 0 |
| | 2 | 0 | 0 | 1 |
| $S_{iu}=6$ | $0,2$ | $(1-p)^2$ | $2p(1-p)$ | $p^2$ |
| | 1 | 0 | 0 | 0 |
| $S_{iu}=5$ | 0 | $1-p$ | $p$ | 0 |
| | 1 | 0 | 0 | 0 |
| | 2 | 0 | $1-p$ | $p$ |
| $S_{iu}=4$ | $0,1,2$ | $1-p$ | 0 | $p$ |
| $S_{iu}=3$ | 0 | 1 | 0 | 0 |
| | 1 | $1/2$ | 0 | $1/2$ |
| | 2 | 0 | 0 | 1 |
| $S_{iu}=2$ | $0,2$ | $1-p$ | 0 | $p$ |
| | 1 | 0 | 0 | 0 |
| $S_{iu}=1$ | 0 | 1 | 0 | 0 |
| | 1 | 0 | 1 | 0 |
| | 2 | 0 | 0 | 1 |

Table 3: $\Pr(G_i|G_u, S_{iu}=k)$

| $S_{u1}$ | $S_{u2}$ | $S_{12}$ |
|---|---|---|
| 9 | 8 | 8 |
| 8 | 8 | 9 |
| 8 | 8 | 8 |
| 8 | 6 | 5 |
| 8 | 5 | 6 |
| 8 | 5 | 5 |
| 5 | 5 | 2 |
| 4 | 3 | 8 |
| 3 | 2 | 5 |

Table 4: Informative 3-way configurations. Note that because individuals 1 and 2 are unordered, we list only one of the corresponding ordered 3-way configurations (e.g. configuration (8, 9, 8) is also informative but has the same probabilities as configuration (9, 8, 8) with individuals 1 and 2 switched). Note that for configuration (8, 8, 7), (7, 7, 7) and (3, 3, 7) $\Pr(G_1, G_2|G_u, S_{u12}) = \Pr(G_1|G_u, S_{u1}) = \Pr(G_2|G_u, S_{u2})$, though we do not seek these configurations in our algorithm.

| | | $G_2$ | | |
|---|---|---|---|---|
| | $G_1$ | 0 | 1 | 2 |
| $G_u = 0$ | 0 | $q^2$ | 0 | 0 |
| | 1 | $pq$ | $pq$ | 0 |
| | 2 | 0 | $p^2$ | 0 |
| $G_u = 1$ | 0 | $q^2/2$ | $q^2/2$ | 0 |
| | 1 | $qp/2$ | $qp$ | $qp/2$ |
| | 2 | 0 | $p^2/2$ | $p^2/2$ |
| $G_u = 2$ | 0 | 0 | $q^2$ | 0 |
| | 1 | 0 | $pq$ | $pq$ |
| | 2 | 0 | 0 | $p^2$ |

Table 5: Configuration ($S_{u1} = 9, S_{u2} = 8, S_{12} = 8$), probabilities $\Pr(G_1, G_2|G_u, S_{u12})$

| | | $G_2$ | | |
|---|---|---|---|---|
| | $G_1$ | 0 | 1 | 2 |
| $G_u = 0$ | 0 | $q^2$ | $pq$ | 0 |
| | 1 | $pq$ | $p^2$ | 0 |
| | 2 | 0 | 0 | 0 |
| $G_u = 1$ | 0 | 0 | $q^2/2$ | $pq/2$ |
| | 1 | $q^2/2$ | $pq$ | $p^2/2$ |
| | 2 | $pq/2$ | $p^2/2$ | 0 |
| $G_u = 2$ | 0 | 0 | 0 | 0 |
| | 1 | 0 | $q^2$ | $pq$ |
| | 2 | 0 | $pq$ | $p^2$ |

Table 6: Configuration ($S_{u1} = 8, S_{u2} = 8, S_{12} = 9$), probabilities $\Pr(G_1, G_2|G_u, S_{u12})$

| | $G_1$ | $G_2$ 0 | 1 | 2 |
|---|---|---|---|---|
| $G_u = 0$ | 0 | $q(q+1)/2$ | $pq/2$ | 0 |
| | 1 | $pq/2$ | $p(p+1)/2$ | 0 |
| | 2 | 0 | 0 | 0 |
| $G_u = 1$ | 0 | $q^2/4$ | $q(p+1)/4$ | 0 |
| | 1 | $q(p+1)/4$ | $(p^2+q^2)/4$ | $p(q+1)/4$ |
| | 2 | 0 | $p(q+1)/4$ | $p^2/4$ |
| $G_u = 2$ | 0 | 0 | 0 | 0 |
| | 1 | 0 | $q(q+1)/2$ | $pq/2$ |
| | 2 | 0 | $pq/2$ | $p(p+1)/2$ |

Table 7: Configuration $(S_{u1} = 8, S_{u2} = 8, S_{12} = 8)$, probabilities $\Pr(G_1, G_2|G_u, S_{u12})$

| | $G_1$ | $G_2$ 0 | 1 | 2 |
|---|---|---|---|---|
| $G_u = 0$ | 0 | $q$ | 0 | 0 |
| | 1 | 0 | 0 | $p$ |
| | 2 | 0 | 0 | 0 |
| $G_u = 1$ | 0 | $q/2$ | 0 | 0 |
| | 1 | $q/2$ | 0 | $p/2$ |
| | 2 | 0 | 0 | $p/2$ |
| $G_u = 2$ | 0 | 0 | 0 | 0 |
| | 1 | $q$ | 0 | 0 |
| | 2 | 0 | 0 | $p$ |

Table 8: Configuration $(S_{u1} = 8, S_{u2} = 6, S_{12} = 5)$, probabilities $\Pr(G_1, G_2|G_u, S_{u12})$

| | $G_1$ | $G_2$ 0 | 1 | 2 |
|---|---|---|---|---|
| $G_u = 0$ | 0 | $q$ | 0 | 0 |
| | 1 | $p$ | 0 | 0 |
| | 2 | 0 | 0 | 0 |
| $G_u = 1$ | 0 | 0 | 0 | $q/2$ |
| | 1 | $q/2$ | 0 | $p/2$ |
| | 2 | $p/2$ | 0 | 0 |
| $G_u = 2$ | 0 | 0 | 0 | 0 |
| | 1 | 0 | 0 | $q$ |
| | 2 | 0 | 0 | $p$ |

Table 9: Configuration $(S_{u1} = 8, S_{u2} = 5, S_{12} = 6)$, probabilities $\Pr(G_1, G_2|G_u, S_{u12})$

| | $G_1$ | $G_2$ 0 | 1 | 2 |
|---|---|---|---|---|
| $G_u = 0$ | 0 | $q$ | 0 | 0 |
| | 1 | $p$ | 0 | 0 |
| | 2 | 0 | 0 | 0 |
| $G_u = 1$ | 0 | $q/2$ | 0 | 0 |
| | 1 | $p/2$ | 0 | $q/2$ |
| | 2 | 0 | 0 | $p/2$ |
| $G_u = 2$ | 0 | 0 | 0 | 0 |
| | 1 | 0 | 0 | $q$ |
| | 2 | 0 | 0 | $p$ |

Table 10: Configuration $(S_{u1} = 8, S_{u2} = 5, S_{12} = 5)$, probabilities $\Pr(G_1, G_2 | G_u, S_{u12})$

| | $G_1$ | $G_2$ 0 | 1 | 2 |
|---|---|---|---|---|
| $G_u = 0$ | 0 | 1 | 0 | 0 |
| | 1 | 0 | 0 | 0 |
| | 2 | 0 | 0 | 0 |
| $G_u = 1$ | 0 | 0 | 0 | 1/2 |
| | 1 | 0 | 0 | 0 |
| | 2 | 1/2 | 0 | 0 |
| $G_u = 2$ | 0 | 0 | 0 | 0 |
| | 1 | 0 | 0 | 0 |
| | 2 | 0 | 0 | 1 |

Table 11: Configuration $(S_{u1} = 5, S_{u2} = 5, S_{12} = 2)$, probabilities $\Pr(G_1, G_2 | G_u, S_{u12})$

| | $G_1$ | $G_2$ 0 | 1 | 2 |
|---|---|---|---|---|
| $G_u = 0$ | 0 | $q^2$ | 0 | 0 |
| | 1 | $pq$ | $pq$ | 0 |
| | 2 | 0 | $p^2$ | 0 |
| $G_u = 1$ | 0 | 0 | 0 | 0 |
| | 1 | 0 | 0 | 0 |
| | 2 | 0 | 0 | 0 |
| $G_u = 2$ | 0 | 0 | $q^2$ | 0 |
| | 1 | 0 | $pq$ | $pq$ |
| | 2 | 0 | 0 | $p^2$ |

Table 12: Configuration $(S_{u1} = 4, S_{u2} = 3, S_{12} = 8)$, probabilities $\Pr(G_1, G_2 | G_u, S_{u12})$

| | $G_1$ | $G_2$ = 0 | 1 | 2 |
|---|---|---|---|---|
| $G_u = 0$ | 0 | $q$ | 0 | 0 |
| | 1 | 0 | 0 | $p$ |
| | 2 | 0 | 0 | 0 |
| $G_u = 1$ | 0 | 0 | 0 | 0 |
| | 1 | 0 | 0 | 0 |
| | 2 | 0 | 0 | 0 |
| $G_u = 2$ | 0 | 0 | 0 | 0 |
| | 1 | $q$ | 0 | 0 |
| | 2 | 0 | 0 | $p$ |

Table 13: Configuration ($S_{u1} = 3, S_{u2} = 2, S_{12} = 5$), probabilities $\Pr(G_1, G_2|G_u, S_{u12})$

| Info | SNP type | MAF | genotype certainty | Number of genotypes (%) | Impute2 | Impute2 & Kinpute |
|---|---|---|---|---|---|---|
| $> 0.4$ | shared | $> 0.02$ | High | 2911525 (59.7) | 0.945 | **0.969** |
| $> 0.4$ | shared | $> 0.02$ | Low | 469987 (9.6) | 0.634 | **0.834** |
| $> 0.4$ | shared | $\leq 0.02$ | High | 387932 (8.0) | 0.869 | **0.931** |
| $> 0.4$ | shared | $\leq 0.02$ | Low | 15460 (0.3) | 0.640 | **0.902** |
| $> 0.4$ | private | $> 0.02$ | High | 201672 (4.1) | 0.927 | **0.960** |
| $> 0.4$ | private | $> 0.02$ | Low | 31896 (0.7) | 0.626 | **0.836** |
| $> 0.4$ | private | $\leq 0.02$ | High | 54182 (1.1) | 0.789 | **0.835** |
| $> 0.4$ | private | $\leq 0.02$ | Low | 3178 (0.1) | 0.691 | **0.824** |
| $< 0.4$ | shared | $> 0.02$ | High | 235481 (4.8) | 0.045 | **0.565** |
| $< 0.4$ | shared | $> 0.02$ | Low | 106855 (2.2) | 0.398 | **0.846** |
| $< 0.4$ | shared | $\leq 0.02$ | High | 284529 (5.8) | 0.000362 | **0.443** |
| $< 0.4$ | shared | $\leq 0.02$ | Low | 8751 (0.2) | 0.0473 | **0.873** |
| $< 0.4$ | private | $> 0.02$ | High | 28684 (0.6) | 0.0240 | **0.671** |
| $< 0.4$ | private | $> 0.02$ | Low | 7988 (0.2) | 0.308 | **0.890** |
| $< 0.4$ | private | $\leq 0.02$ | High | 123111 (2.5) | 0.00295 | **0.158** |
| $< 0.4$ | private | $\leq 0.02$ | Low | 5913 (0.1) | 0.0843 | **0.654** |

Table 14: Heterozygote sensitivity

| Info | SNP type | MAF | genotype certainty | Number of genotypes (%) | Impute2 | Impute2 & Kinpute |
|---|---|---|---|---|---|---|
| $> 0.4$ | shared | $> 0.02$ | High | 2911525 (59.7) | **0.953** | 0.950 |
| $> 0.4$ | shared | $> 0.02$ | Low | 469987 (9.6) | 0.590 | **0.766** |
| $> 0.4$ | shared | $\leq 0.02$ | High | 387932 (8.0) | 0.961 | **0.965** |
| $> 0.4$ | shared | $\leq 0.02$ | Low | 15460 (0.3) | 0.758 | **0.909** |
| $> 0.4$ | private | $> 0.02$ | High | 201672 (4.1) | **0.934** | 0.932 |
| $> 0.4$ | private | $> 0.02$ | Low | 31896 (0.7) | 0.579 | **0.752** |
| $> 0.4$ | private | $\leq 0.02$ | High | 54182 (1.1) | 0.670 | **0.755** |
| $> 0.4$ | private | $\leq 0.02$ | Low | 3178 (0.1) | 0.330 | **0.630** |
| $< 0.4$ | shared | $> 0.02$ | High | 235481 (4.8) | 0.531 | **0.761** |
| $< 0.4$ | shared | $> 0.02$ | Low | 106855 (2.2) | 0.437 | **0.741** |
| $< 0.4$ | shared | $\leq 0.02$ | High | 284529 (5.8) | 0.429 | **0.923** |
| $< 0.4$ | shared | $\leq 0.02$ | Low | 8751 (0.2) | 0.502 | **0.892** |
| $< 0.4$ | private | $> 0.02$ | High | 28684 (0.6) | 0.493 | **0.844** |
| $< 0.4$ | private | $> 0.02$ | Low | 7988 (0.2) | 0.404 | **0.733** |
| $< 0.4$ | private | $\leq 0.02$ | High | 123111 (2.5) | 0.211 | **0.585** |
| $< 0.4$ | private | $\leq 0.02$ | Low | 5913 (0.1) | 0.109 | **0.391** |

Table 15: Heterozygote positive predictive value

| Info | SNP type | MAF | genotype certainty | Number of genotypes (%) | Impute2 | Impute2 & Kinpute |
|---|---|---|---|---|---|---|
| $> 0.4$ | shared | $> 0.02$ | High | 2911525 (59.7) | 0.964 | **0.972** |
| $> 0.4$ | shared | $> 0.02$ | Low | 469987 (9.6) | 0.611 | **0.810** |
| $> 0.4$ | shared | $\leq 0.02$ | High | 387932 (8.0) | 0.987 | **0.992** |
| $> 0.4$ | shared | $\leq 0.02$ | Low | 15460 (0.3) | 0.661 | **0.888** |
| $> 0.4$ | private | $> 0.02$ | High | 201672 (4.1) | 0.966 | **0.973** |
| $> 0.4$ | private | $> 0.02$ | Low | 31896 (0.7) | 0.631 | **0.810** |
| $> 0.4$ | private | $\leq 0.02$ | High | 54182 (1.1) | 0.983 | **0.987** |
| $> 0.4$ | private | $\leq 0.02$ | Low | 3178 (0.1) | 0.682 | **0.879** |
| $< 0.4$ | shared | $> 0.02$ | High | 235481 (4.8) | 0.933 | **0.961** |
| $< 0.4$ | shared | $> 0.02$ | Low | 106855 (2.2) | 0.569 | **0.838** |
| $< 0.4$ | shared | $\leq 0.02$ | High | 284529 (5.8) | 0.970 | **0.982** |
| $< 0.4$ | shared | $\leq 0.02$ | Low | 8751 (0.2) | 0.612 | **0.912** |
| $< 0.4$ | private | $> 0.02$ | High | 28684 (0.6) | 0.945 | **0.975** |
| $< 0.4$ | private | $> 0.02$ | Low | 7988 (0.2) | 0.615 | **0.862** |
| $< 0.4$ | private | $\leq 0.02$ | High | 123111 (2.5) | **0.989** | **0.989** |
| $< 0.4$ | private | $\leq 0.02$ | Low | 5913 (0.1) | 0.901 | **0.919** |

Table 16: Concordance rates
